# Loss of Shh signaling in the neocortex reveals heterogeneous cell recovery responses from distinct oligodendrocyte populations

**DOI:** 10.1101/547968

**Authors:** Caitlin C Winkler, Santos J Franco

**Author notes:** Corresponding Author: University of Colorado - AMC, 12800 E. 19.

## Abstract

The majority of oligodendrocytes in the neocortex originate from neural progenitors that reside in the dorsal forebrain. We recently showed that Sonic Hedgehog (Shh) signaling in these dorsal progenitors is required to produce normal numbers of neocortical oligodendrocytes during embryonic development. Conditional deletion of the Shh signaling effector, *Smo*, in dorsal progenitors caused a dramatic reduction in oligodendrocyte numbers in the embryonic neocortex. In the current study, we show that the depleted oligodendrocyte lineage in *Smo* conditional mutants is able to recover to control numbers over time. This eventual recovery is achieved in part by expansion of the ventrally-derived wild-type lineage that normally makes up a minority of the total oligodendrocyte population. However, we find that the remaining dorsally-derived mutant cells also increase in numbers over time to contribute equally to the recovery of the total population. Additionally, we found that the ways in which the dorsal and ventral sources cooperate to achieve recovery is different for distinct subclasses of oligodendrocyte-lineage cells. Oligodendrocyte precursor cells (OPCs) in the neocortical white matter recover completely by expansion of the remaining dorsally-derived *Smo* mutant cells. On the other hand, mature oligodendrocytes in the white and gray matter recover through an equal contribution from dorsal mutant and ventral wild-type lineages. Interestingly, the only population that did not make a full recovery was OPCs in the gray matter. We find that gray matter OPCs are less proliferative in *Smo* cKO mutants compared to controls, which may explain their inability to fully recover. These studies shed light on the nature of competition and cooperation between dorsal and ventral sources of oligodendrocytes in the developing neocortex. Furthermore, our data point towards differential use of Shh signaling in the development of distinct subclasses of the oligodendrocyte lineage.

## Introduction

Development of neural circuits in the neocortex requires coordinated production of many distinct cell types with specialized functions, including different types of neurons and glia. Oligodendrocytes are glial cells that are responsible for myelinating neuronal axons to increase electrical conduction velocity and provide trophic support. In addition, oligodendrocytes and their precursor cells (OPCs) can form synapse-like connections with neurons to engage in activity-dependent neuron-glia signaling. Thus, oligodendrocytes play a major role in creating the structural and electrical landscape in the neocortex. Elucidating the cellular and molecular mechanisms of oligodendrocyte development is therefore critical for understanding neural circuit formation and function in the neocortex.

During embryonic development, neocortical oligodendrocytes are generated from oligodendrocyte precursor cells (OPCs) that arise in three distinct waves. The first two waves come from the ganglionic eminences in the ventral forebrain and the third wave is produced locally from neural progenitors in the dorsal forebrain (Kessaris et al., 2005; Richardson et al., 2006). OPCs derived from ventral sources initially infiltrate and populate the developing neocortex, but are quickly outnumbered by dorsally-derived OPCs that ultimately contribute 80-90% of all oligodendrocytes and OPCs in the mature neocortex (Kessaris et al., 2005; Tripathi et al., 2011; Winkler et al., 2018). The function of this ventral-to-dorsal switch is not yet known, nor are the mechanisms driving the exact ratios between the different sources. In fact, several studies indicate that the different oligodendrocyte lineages are molecularly and physiologically very similar once they reach the neocortex (Marques et al., 2018; Tripathi et al., 2011), and are even functionally interchangeable (Kessaris et al., 2005). However, other studies have demonstrated functional diversity of OPCs and oligodendrocytes in the spinal cord and neocortex based on their dorsal or ventral origins (Crawford et al., 2016; Tripathi et al., 2011; Zhu et al., 2011). There also appear to be regional differences among oligodendrocytes based on their location in the white matter or gray matter (Dimou et al., 2008; Hill et al., 2013; Kang et al., 2010; Viganò et al., 2013; Young et al., 2013). Additionally, several studies have demonstrated that distinct populations of oligodendrocytes and OPCs require different morphogens and utilize different signaling pathways. Whereas some oligodendrocytes require Sonic hedgehog (Shh) signaling (Fuccillo, 2004; Spassky et al., 2001; Tekki-Kessaris et al., 2001; Tong et al., 2015), others are able to differentiate and proliferate via a Shh-independent mechanism (Cai et al., 2005; Chandran et al., 2003; Fogarty et al., 2005; Hashimoto et al., 2017; Nery et al., 2001; Vallstedt et al., 2005). Furthermore, specific levels of Shh signaling have recently been shown to drive differential fate determination between oligodendrocytes and NG2^+^ OPCs (Ravanelli et al., 2018). Together, these studies have provided evidence for an increasingly complex heterogeneity amongst the oligodendroglial lineage based on developmental origin, regional location and molecular signaling pathways.

In this study, we investigated the interplay between the dorsal and ventral oligodendrocyte sources in the developing mouse neocortex, and the roles that Shh signaling plays in different oligodendrocyte subclasses within the the dorsal lineage. Previously, we used conditional knockout approaches to show that highly regulated Sonic hedgehog (Shh) signaling from multiple embryonic sources is necessary for normal oligodendrocyte production from dorsal forebrain progenitors (Winkler et al., 2018). When we used an *Emx1-Cre* driver line to conditionally knock out the obligate Shh pathway component, *Smoothened* (*Smo*), in dorsal forebrain progenitors, the neocortical oligodendrocyte population was severely diminished by embryonic day (E) 17.5. Importantly, however, a small population of Olig2^+^ oligodendrocyte-lineage cells was preserved in the mutants. This remaining population consisted of a mix of dorsally-derived *Smo* mutant cells and ventrally-derived *Smo* wild-type cells, which provided us with an experimental paradigm to study how these two lineages coordinately responded during subsequent postnatal brain development. Here, we find that the overall numbers of OPCs and oligodendrocytes eventually recover to normal levels in the mature neocortex, despite being dramatically reduced after loss of Shh signaling embryonically. Furthermore, we show that the dorsally-derived *Smo* mutant cells and the ventrally-derived wild-type cells both contribute significantly to the overall recovery of the Olig2^+^ cell population in *Emx1;Smo* mutants. Interestingly, we find that different classes of oligodendrocyte-lineage cells use distinct dorsal:ventral ratios to recover, depending on their regional location in the gray or white matter and their maturational state as OPCs or mature oligodendrocytes. Finally, we used this experimental paradigm to begin to understand differential requirements of Shh signaling in these distinct subclasses within the dorsal oligodendroglial lineage. Specifically, we found that dorsally-derived OPCs in the white matter exhibited the most robust recovery after loss of Shh signaling, whereas OPCs in the gray matter could never fully recover. Together, these studies provide insights into the developmental dynamics and heterogeneity of neocortical oligodendrocytes, with respect to several emerging properties of oligodendrocyte diversity: developmental origin (dorsally-derived vs. ventrally-derived), regional niche (WM vs. GM), maturational states (OPCs vs. OLs), and signaling pathways (Shh-dependent vs. –independent).

## MATERIALS AND METHODS

### Mice

The following mice were obtained from The Jackson Laboratory: *Emx1-Cre* (*B6.129S2-Emx1*^*tm1(cre)Krj*^*/J*, stock no. 005628); *R26-NZG* [*FVB.Cg-Gt(ROSA)26Sor*^*tm1(CAG-lacZ,-EGFP)Glh*^*/J*, stock no. 012429]; *Smo-floxed* (*Smo*^*tm2Amc*^*/J*, stock no. 004526). *Emx1-Cre* mice were crossed with *R26-NZG* mice to generate double heterozygous animals. *Emx1*^*+/Cre*^;*R26*^+*/NZG*^ double heterozygotes were then crossed with *Smo-floxed* animals to generate *Emx1*^+*/Cre*^*;R26*^+*/NZG*^*;Smo*^+*/fl*^ triple heterozygotes. The triple heterozygous animals were mated back to *Smo*^*fl/fl*^ animals to generate conditional knock-out mice of the informative genotype *Emx1*^+*/Cre*^*;R26*^+*/NZG*^*;Smo*^*fl/fl*^ (*Smo* cKO). *Emx1*^+*/Cre*^*;R26*^+*/NZG*^*;Smo*^+*/*+^ littermates were used as wild-type controls(*Smo* control). Animals were maintained according to the guidelines from the Institutional Animal Care and Use Committee of the University of Colorado, Denver, or the University of California, San Francisco. Male and female mice were used equally throughout our experiments.

### Tissue preparation

Embryonic brains were fixed in 4% paraformaldehyde (PFA) for 1 h at room temperature (RT). Postnatal day 4 (P4) brains were fixed in 4% PFA overnight at 4 degrees C. All other postnatal mice were transcardially perfused with 4% PFA and brains postfixed in 4% PFA for 1 h at RT. For immunohistochemistry, brains were sectioned coronally at 50–100 µm with a vibrating microtome. For RNAscope® mRNA in situ hybridization, brains were cryoprotected in 30% sucrose at 4 degrees C overnight and sectioned on a cryostat at 12 µm.

### Immunohistochemistry

Free-floating vibratome sections were blocked with 10% donkey serum and 0.2% Triton-X in 1X PBS for 2 h at RT. After 2 h, the blocking solution was removed and sections were incubated with primary antibodies in 10% donkey serum in 1X PBS for 1 h at RT, and then washed at RT with 1X PBS three times for 5 min each. After washing, sections were incubated with secondary antibodies in 10% donkey serum in 1X PBS for 1 h at RT, and then washed again with 1X PBS three times for 5 min each. Sections were mounted on slides with ProLong Diamond Antifade Mountant (Invitrogen). Images were captured using a LSM780 Zeiss laser-scanning confocal microscope. Antibodies used for immunostaining were as follows: rabbit anti-Olig2 (1:500; Millipore, RRID:AB_2299035), rat anti-PDGFRα (1:500; BD Pharmingen; RRID:AB_397117), chicken anti-β-gal (1:2000; Abcam; RRID:AB_307210), mouse anti-Ki67 (1:250; BD Pharmingen; RRID:AB_393778). Donkey secondary antibodies conjugated to Alexa Fluor 488, Rhodamine Red-X, or Alexa Fluor 647 were purchased from Jackson ImmunoResearch and used at 1:500.

### RNAscope® mRNA in situ hybridization combined with immunohistochemistry

mRNA in situ hybridization for *Smo* was performed using the RNAscope® Multiplex Fluorescent Reagent Kit v2 (Advanced Cell Diagnostics) with a *Smo* probe (RNAscope® Probe – Mm-*Smo*; Advanced Cell Diagnostics, catalog no. 318411). The assay was performed in accordance with the manufacturer’s instructions, then followed by immunohistochemistry for Olig2 and βGal. After completing the RNAscope® assay, sections were washed 2x for 2 min each in TBS pH 7.6 plus 1% bovine serum albumin (TBS-B). Sections were then blocked in 10% normal donkey serum in TBS-B for 30 min at RT. Primary antibodies (same as above) were diluted in TBS-B and incubated on the sections for 2 hours at RT, followed by washing 3x for 5 min each in TBS plus 0.005 Tween® 20 (TBS-T). Horseradish peroxidase-conjugated secondary antibodies were diluted 1:250 in TBS-B. Secondary antibodies were: donkey anti-chicken (RRID: AB_10015283) and donkey anti-rabbit (RRID: AB_10015282) from Jackson ImmunoResearch. Sections were first incubated in anti-chicken secondary for 30 min at RT, followed by washing 3x for 5 min each in TBS-T. Slides were then incubated with Opal™ 570 reagent (1:1000; PerkinElmer) for 10 min at RT. Slides were rinsed 3x for 2 min each in TBS-T, followed by quenching of the peroxidase activity using HRP blocker from the RNAscope kit, 30 min at 40 degrees C. The secondary and Opal™ steps were then repeated using the anti-rabbit secondary and Opal™ 650 reagent (1:1000; PerkinElmer). Sections were then washed 3x for 2 min each in TBS-T and counterstained with DAPI to label nuclei.

### Experimental design and statistical analysis

For all immunostainings, ≥3 histological sections at three distinct rostrocaudal levels from each of three different animals at each time point (nine sections total for each condition) were analyzed. Analysis was restricted to the mediolateral part of the neocortex comprising primarily the presumptive somatosensory cortex in embryonic brains and the somatosensory cortex in postnatal brains. Biological replicates are individual animals. Confocal, single-plane optical sections were used for quantification. Cells were analyzed in contiguous columns spanning the entire neocortex from ventricle to pial surface. Percentages of marker-positive cells were quantified from these columns. Absolute numbers of marker-positive cells were also quantified from these columns and then divided by the area of a column to get cell density (cells/mm^2^). Normalized cell counts are cell densities normalized to control counts. Values in graphs are mean ± SEM for biological replicates. For independent two-group experiments (e.g., WT vs cKO), an unpaired two-tailed Student’s *t* test was used to determine statistical significance. For analysis involving ≥3 independent groups, a one-way ANOVA was used followed by Tukey’s *post hoc* test. Values were considered statistically significant at *p* ≤ 0.05.

## RESULTS

### Postnatal recovery of neocortical Olig2^+^ cells after embryonic depletion in *Smo* conditional knockout brains

We previously showed that the Sonic Hedgehog (Shh) signaling pathway is required in dorsal forebrain progenitors to generate normal numbers of oligodendrocyte precursor cells (OPCs) in the embryonic neocortex (Winkler et al., 2018). Specifically, we used the *Emx1-Cre* driver to conditionally knock out the obligate Shh pathway component, *Smoothened* (*Smo*), in dorsal forebrain progenitors early in embryonic development (Fig. 1A). Conditional knockout (cKO) of *Smo* in dorsal progenitors led to a dramatic reduction in total Olig2^+^ cells in the neocortex by embryonic day 17.5 (E17.5) (Fig. 1B). Importantly, a small proportion of Olig2^+^ cells remained in the *Emx1;Smo* mutants at E17.5 (Fig. 1B-C). Because OPCs are proliferative and remain so during postnatal development, we wondered if this loss of Olig2^+^ cells persisted in the mutant brain or if the oligodendrocyte lineage could eventually recover. We allowed the *Emx1;Smo* mutants to develop until postnatal day 30 (P30), and stained brain sections for Olig2 to label OPCs and mature oligodendrocytes. We found that by P30 the total Olig2^+^ population was largely recovered in the *Smo* cKO mutants (Figure 1D), coming back to ∼75% of control wild-type (WT) littermates (Fig. 1E). These data indicated a robust recovery of the oligodendrocyte lineage in the *Smo* cKO neocortex, despite starting with significantly depleted numbers at the onset of oligodendrogenesis.

**Figure 1.**
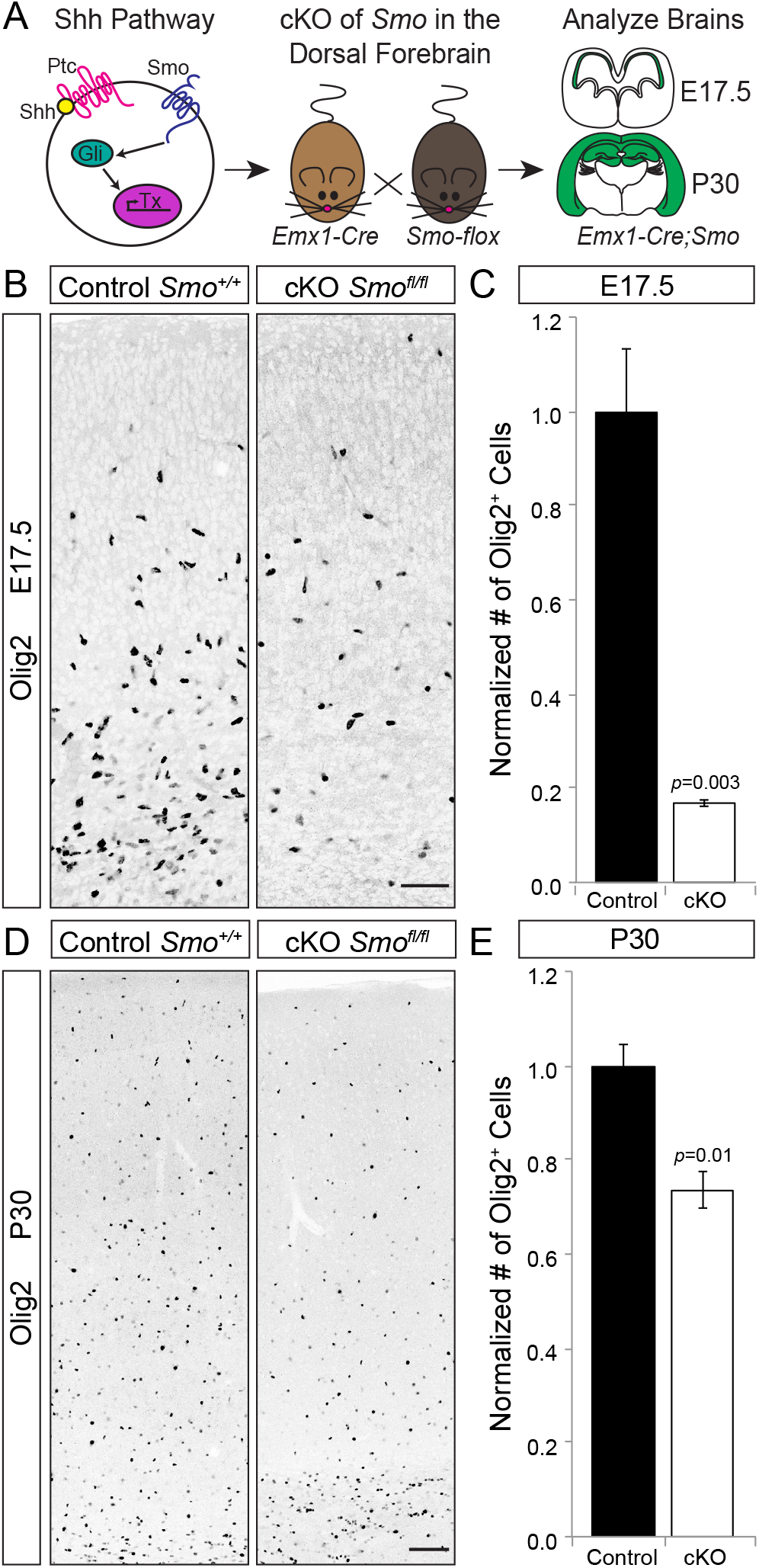
Oligodendrocyte-lineage cells in *Smo* cKO mutants largely recover by P30 after dramatic reduction at E17.5. ***A*,** Schematic of experimental design, modified from Winkler et al., 2018. Shh binding to its receptor Patched (Ptc) allows the essential downstream effector Smo to activate Gli transcription factors and initiate downstream transcription. *Smo* was knocked out in just the dorsal progenitor population by crossing *Smo-flox mice* to *Emx1-Cre* mice. Brains were analyzed at E17.5 and P30 for the effect on neo-cortical oligodendrocytes, identified by immunostaining for Olig2. ***B*,** Representative overview images of the presumptive somatosensory cortex at E17.5, showing loss of oligodendrocyte-lineage cells (Olig2^+^) in *Emx-1;Smo* mutant brains (right) compared with controls (left). ***C*,** Quantification of Olig2^+^ cell loss in the E17.5 neocortex. Graph is number of Olig2^+^ cells in a standard-size area, normalized to controls (±SEM among biological replicates). ***D*,** Representative overview images of the adult somatosensory cortex at P30, showing recovery of oligodendrocyte-lineage cells in *Emx1;Smo* mutant brains (right) compared with controls (left). ***E*,** Quantification of oligodendrocyte-lineage cells in the P30 somatosensory cortex, showing that *Emx1;Smo* mutants had largely recovered relative to controls. Graph as in (C) (±SEM among biological replicates). Scale bar, 100 μm.

Interestingly, our previous study indicated that the remaining Olig2^+^ cells in the E17.5 *Emx1;Smo* cKO neocortex were a mix of dorsally-derived *Smo* mutant cells and ventrally-derived *Smo* wild-type cells (Winkler et al., 2018). These two populations can be distinguished by their expression (dorsal cKO) or absence (ventral WT) of a Cre-dependent βGal transgene (Fig. 2A). We used this strategy to quantify the percentage of Olig2^+^ cells that were dorsally-derived (βGal^+^) in *Smo* control and cKO neocortex at E17.5 (Fig. 2B-C). Consistent with our previous study (Winkler et al., 2018), we found that ∼90% of Olig2^+^ cells in the E17.5 control neocortex were dorsally-derived (Fig. 2C). In the *Smo* cKO mutants, however, the dorsal population dropped significantly to only 27% of the total Olig2^+^ cells (Fig. 2C). We confirmed that the remaining dorsally-derived cells were in fact *Smo* knockouts by combining RNAscope in situ hybridization for *Smo* mRNA with immunohistochemistry for Olig2 and βGal protein (Fig. 2A). Indeed, we never detected *Smo* mRNA in any Olig2^+^βGal^+^ double-positive cells in the neocortex (Fig. 2D). In contrast, *Smo* mRNA was readily detectable in the majority of βGal^−^ cells, including the ventrally-derived Olig2^+^βGal^−^ population (Fig. 2D). Together, these data indicate that loss of Shh signaling to dorsal progenitors led to a significant reduction in oligodendrocyte-lineage cells in the embryonic neocortex, and that the small numbers of remaining Olig2^+^ cells comprise both *Smo* mutant and wild-type cells from dorsal and ventral sources, respectively. Therefore, this gave us an experimental paradigm to study which population, dorsally-derived mutants or ventrally-derived controls, was responsible for the eventual recovery of Olig2^+^ cells in the mature *Smo* cKO neocortex.

**Figure 2.**
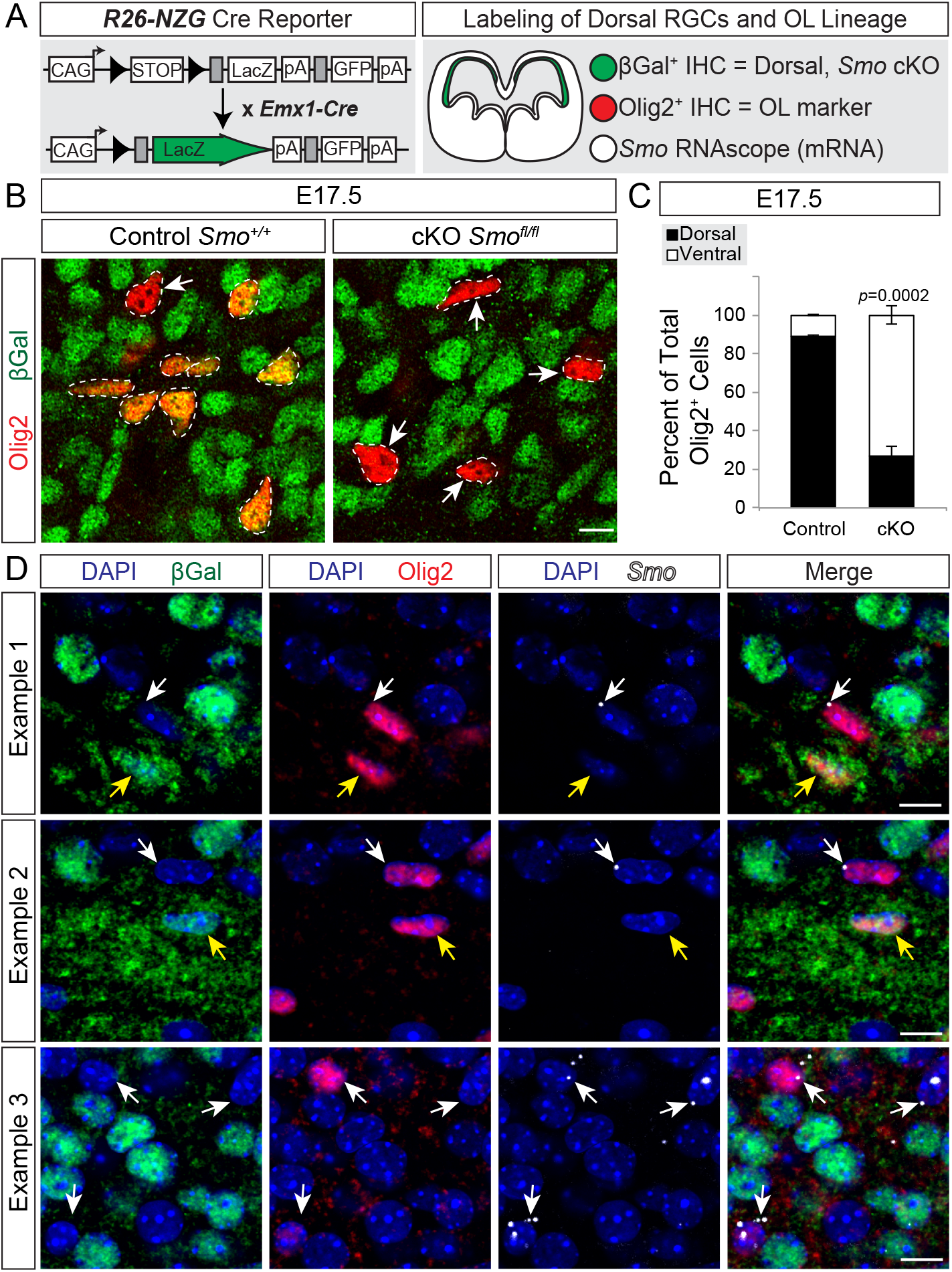
The remaining oligodendrocyte-lineage cells in *Emx1;Smo* mutants are a mix of dorsally-derived *Smo* mutants and ventrally-derived *Smo* wild-type cells. ***A*,** Schematic of the genetic lineage-tracing approach to distinguish dorsally derived cells in the somatosensory cortex from ventrally derived cells. Modified from Winkler et al., 2018. *Emx1-Cre;Smo-flox* mice were crossed to *R26-NZG* reporter mice to permanently label all Emx1^+^ progenitors and their offspring (βGal^+^) and identify oligodendrocyte lineage cells by immunostaining for Olig2. Immunohistochemistry for these two markers was combined with RNAscope in situ hybridization for *Smo* mRNA. ***B*,** Examples of Olig2^+^ cells (red, outlined) in E17.5 *Emx1;Smo* controls (left) and mutants (right) that are either βGal^+^ (green) and therefore dorsally derived, or βGal^−^ and ventrally-derived (white arrows). ***C*,** Graph of the average percentage (±SEM among biological replicates) of Olig2^+^ cells that were dorsally derived (βGal^+^) versus ventrally derived (βGal^−^) in the E17.5 neocortex. ***D*,** Immunohistochemistry as in (B), combined with RNAscope in situ hybridization for Smo mRNA. RNAscope signal for *Smo* mRNA appears as small dots within positive cells (white). Scale bars: (B), 20 μm; (D), 10 μm.

### Repopulation of the oligodendrocyte lineage is achieved by recovery of dorsally-derived mutant cells and expansion of ventrally-derived wild-type cells

Previous studies showed that dorsal and ventral sources of neocortical OPCs can compensate for the loss of one another, ultimately achieving normal OPC numbers in the absence of one source (Kessaris et al., 2005). Additionally, it has been proposed that dorsal and ventral sources of OPCs might compete for mitogenic or survival factors in the developing neocortex (Kessaris et al., 2005; Richardson et al., 2006), similar to what is observed in the spinal cord (Calver et al., 1998; van Heyningen et al., 2001). Because ventrally-derived OPCs outnumbered their dorsal counterparts in the E17.5 *Smo* Cko neocortex by more than 2:1 (Fig. 2D), we reasoned that the ventral OPCs might compensate during the postnatal recovery phase and outcompete the dorsal OPCs by P30. To test this, we quantified the percentage of Olig2^+^ cells that were dorsally-derived (βGal^+^) and ventrally-derived (βGal^−^) in *Smo* control and cKO neocortex at P30 (Fig. 3A-B). Indeed, we found that dorsal cells made up ∼95% of the Olig2^+^ population in control brains at P30, whereas in the *Smo* cKO neocortex the Olig2^+^ population comprised 43% dorsal and 57% ventral cells (Fig. 3B). This shifted dorsal:ventral ratio was accompanied by an ∼8-fold increase in the number of ventrally-derived Olig2^+^ cells in *Smo* cKO brains compared to controls (Fig. 3C). Thus, despite being at normal control levels in the E17.5 mutant brain (Fig. 3C), ventrally-derived OPCs appeared to subsequently expand by P30 to partially compensate for loss of Olig2^+^ cells in the *Smo* cKO neocortex. Interestingly, however, we noticed that the dorsally-derived Olig2^+^ cells also recovered substantially between E17.5 and P30 (Fig. 3D). At E17.5, dorsally-derived Olig2^+^ cells were only at ∼5% of control numbers, but by P30 they had recovered to ∼33% of control levels (Fig. 3D). This dorsal recovery was reflected in the fact that the dorsal:ventral ratio had shifted from nearly 1:2 at E17.5 to closer to 1:1 by P30 (Fig. 3B). When we analyzed the recovery of total Olig2^+^ cells in the P30 *Smo* cKO mutant (Fig. 1E) in the context of relative levels of the ventral (Fig. 3C) and dorsal (Fig. 3D) populations, we could see that that the partial recovery at P30 was accomplished by a significant expansion of both ventrally-derived and dorsally-derived Olig2^+^ cells (Fig. 3E). Taken together, these data confirm that ventrally-derived OPCs can indeed partially compensate for loss of dorsally-derived OPCs, but indicate that ventral cells do not outcompete their dorsal counterparts despite outnumbering them. Furthermore, these data show that even in the absence of Shh signaling a small population of dorsally-derived OPCs can expand to populate the postnatal neocortex.

**Figure 3.**
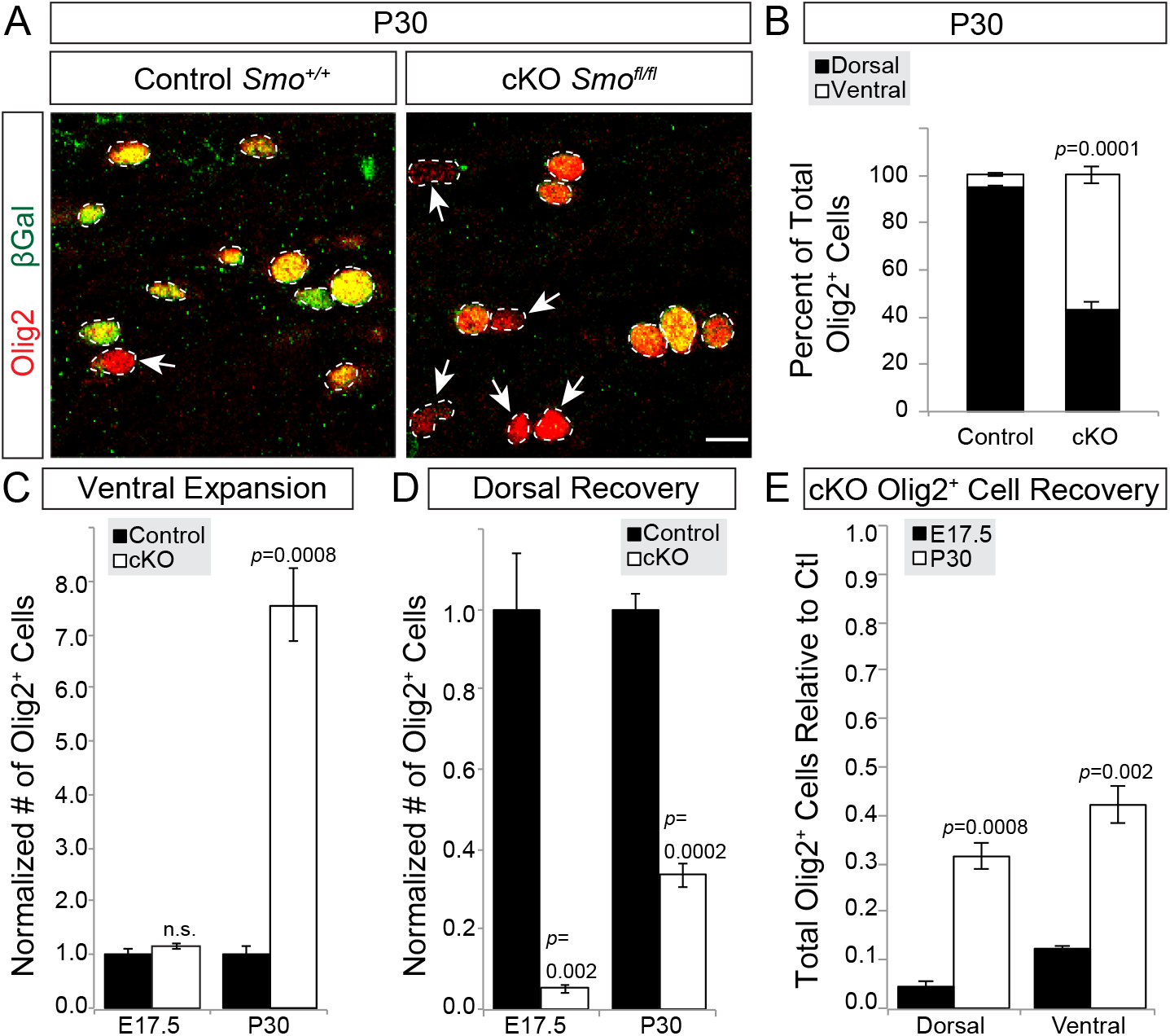
Oligodendrocyte-lineage cells in *Emx1;Smo* mutants recover due to expansion of both dorsal and ventral sources. ***A*,** Examples of Olig2^+^ (red, outlined) cells in *Emx1;Smo* controls (left) and mutants (right) that are either βGal^+^ (green) and therefore dorsally derived, or βGal^−^ and ventrally-derived (white arrows) at P30. ***B*,** Graph of the average percentage (±SEM among biological replicates) of Olig2^+^ cells that were dorsally derived (βGal^+^) versus ventrally derived (βGal^−^) in the P30 neocortex. ***C-D*,** Quantification of Olig2^+^ cells in *Emx1;Smo* controls and mutants at E17.5 and P30. Ventrally-derived Olig2^+^ cells (C) in the mutants are at normal levels at E17.5, but are significantly expanded compared to controls at P30. Dorsally-derived Olig2^+^ cells (D) are greatly depleted in mutants at E17.5, but recover significantly by P30. Graphs are number of Olig2^+^ cells in a standard-size area, normalized to age-matched controls (±SEM among biological replicates). ***E*,** Quantification of the increases over time for the dorsal (βGal^+^Olig2^+^) and ventral (βGal^−^Olig2^+^) populations in *Emx1;Smo* mutants, showing that both population increase between E17.5 and P30. Graph represents the levels of dorsal and ventral Olig2^+^ cells in the *Smo* cKO mutants relative to controls at each age. Levels are calculated from total normalized Olig2^+^ cell numbers at E17.5 (from Fig. 1C) and and P30 (from Fig. 1E), multiplied by the percent dorsal:ventral contribution at E17.5 (from Fig. 2C) or P30 (from Fig. 3B). Scale bar, 20 μm.

We next wanted to gain further insight into this partial recovery of the oligodendrocyte lineage by examining the temporal progression of the Olig2^+^ cell expansion from E17.5 to P30. We analyzed *Smo* control and cKO animals at P4 and P10, two intermediate time points between E17.5 and P30 when the neocortex and its resident oligodendrocyte populations are rapidly expanding. We found that as early as P4 the total Olig2^+^ population in *Emx1;Smo* mutants had already substantially recovered toward control numbers (Figure 4A). Interestingly, by P4 the percent recovery of the total Olig2^+^ population had already reached a level similar to P10 and P30 (Fig. 4A). We observed the same temporal pattern when we separated the dorsal and ventral populations. Dorsally-derived Olig2^+^ cells increased substantially between E17.5 and P4, but then failed to significantly recover any further at P10 or P30 (Fig. 4B). Similarly, ventrally-derived Olig2^+^ cells were greatly expanded relative to controls by P4, but this relative increase plateaued between P4 and P30 (Fig. 4C). These behaviors were reflected in the dorsal:ventral ratios over time, which did not change significantly between P4, P10 and P30 (Fig. 4D, one-way ANOVA: *F*_(2,6)_=0.868, *p*=0.47). Together, these data indicate that expansion of the remaining Olig2^+^ cells in the *Smo* cKO neocortex appears to occur in a short perinatal window, and subsequently remains steady throughout later postnatal ages.

**Figure 4.**
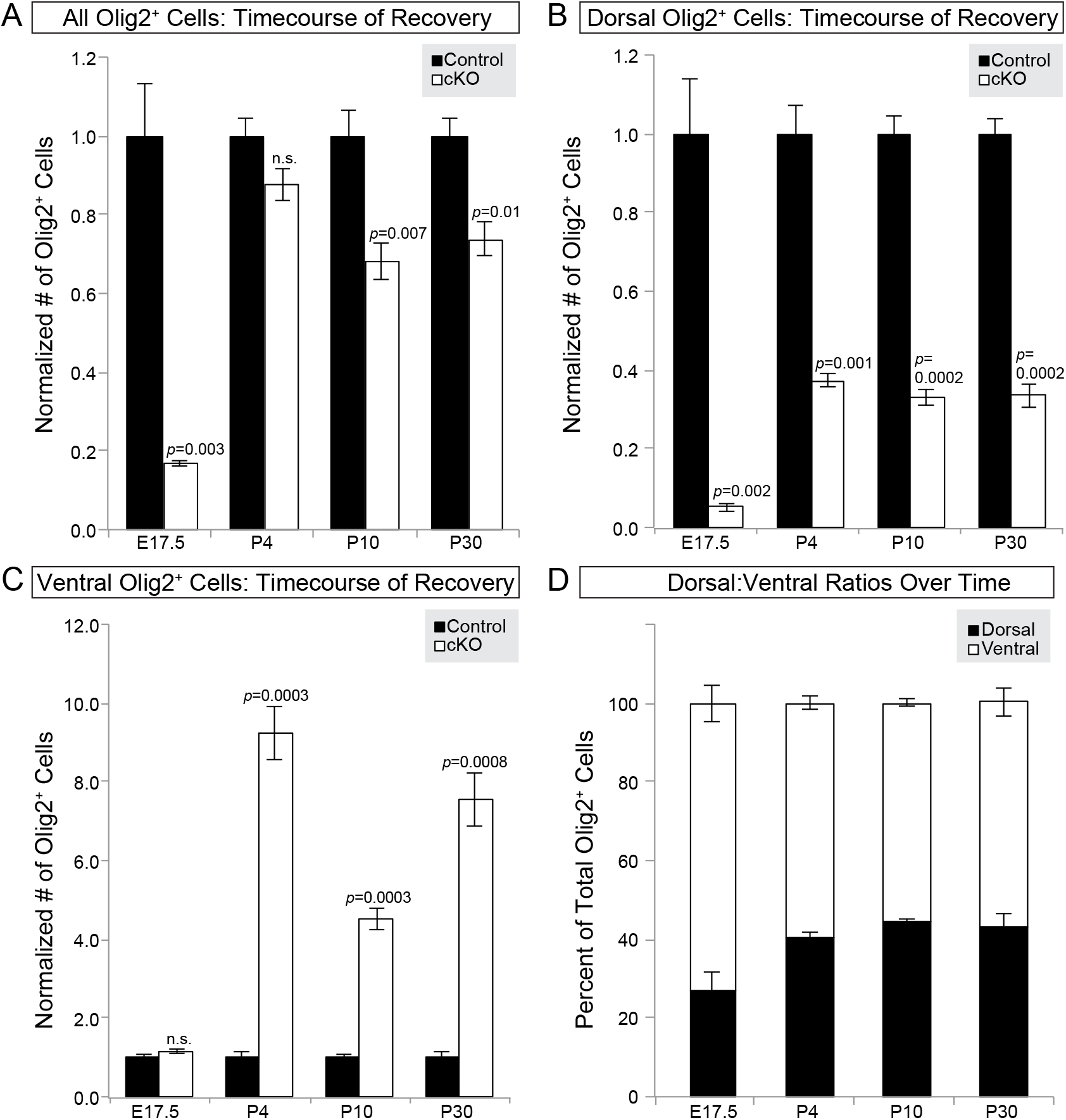
Expansion of the dorsally-derived oligodendrocyte lineage peaks by P4, and recovery of the total oligodendrocyte population stabilizes by P10 in *Emx1;Smo* mutants. ***A*,** The total Olig2^+^ cell population is dramatically reduced in the E17.5 mutant neocortex relative to controls. However, total Olig2^+^ cells largely recover by P4 and stabilize by P10. Graph is number of Olig2^+^ cells in a standard-size area, normalized to controls (±SEM among biological replicates). ***B*,** The total number of dorsally-derived oligodendrocyte lineage cells (βGal^+^Olig2^+^) in the mutant neocortex sharply increases from E17.5 to P4, then plateaus through later timepoints. Graph as in (A). ***C*,** The total number of ventrally-derived oligodendrocyte lineage cells (βGal^−^Olig2^+^) in the mutant neocortex dramatically expands from E17.5 to P4 and continues to exceed control numbers through later timepoints. Graph as in (C). ***D*,** Quantification of the average percentage (±SEM among biological replicates) of Olig2^+^ cells in the mutant neocortex that were dorsally derived (βGal^+^) versus ventrally derived (βGal^−^) at different timepoints. The overall contribution by the dorsally-derived oligodendrocyte lineage increases between E17.5 and P4, then stabilizes across P10 and P30 time-points. Quantification for E17.5 and P30 timepoints were re-graphed from Figures 1 and 2.

### Specific subpopulations of the oligodendrocyte lineage are more susceptible to loss of Shh signaling

Although the overall levels of Olig2^+^ cells returned to normal in the *Smo* cKO neocortex, we wondered if there were specific categories of oligodendrocyte-lineage cells that were more affected than others. For example, were there certain populations in which the dorsal oligodendrocyte lineage recovered more robustly, while others required a greater compensation from the ventral lineage? If so, this might provide some insight into which populations are more reliant on Shh signaling. To test this, we analyzed *Smo* control and cKO brains at P30 for differences within two major categories of the oligodendrocyte lineage: 1) location in the gray matter (GM) vs. white matter (WM) (Fig. 5A), and 2) OPCs vs. mature oligodendrocytes (OLs) (Fig. 5B). We focused on the dorsally-derived (βGal^+^) cells to determine if loss of Shh signaling specifically affected any of these populations. We found that dorsally-derived Olig2^+^ cells (OPCs plus OLs) recovered similarly in the gray matter (38%) and white matter (28%) of *Smo* cKO mutants (*p*=0.18) (Fig. 5B). Likewise, dorsally-derived OPCs and mature OLs recovered to similar levels as each other (43% and 32%, respectively. *p*=0.24) (Fig. 5C). We next analyzed the four populations separately: GM OLs, WM OLs, GM OPCs, and WM OPCs. Here again, we found that mature OLs recovered to similar levels regardless of their location in the GM (34%) or WM (29%) (Fig 5D). However, dorsally-derived WM OPCs in *Smo* cKO mutants did not recover to the same extent as their GM OPC counterparts (19% and 47%, respectively) (Fig. 5D). A one-way ANOVA indicated a significant difference between the percent recovery among the four different categories of Olig2^+^ cells (*F*_(3,8)_=4.704, *p*=0.0355). A *post-hoc* Tukey’s test revealed a significant difference in the percent recovery between WM and GM OPCs (*p*=0.026). Taken together, these data seemed to suggest that WM OPCs were the slowest population to recover at P30 after loss of Shh signaling during embryonic development.

**Figure 5.**
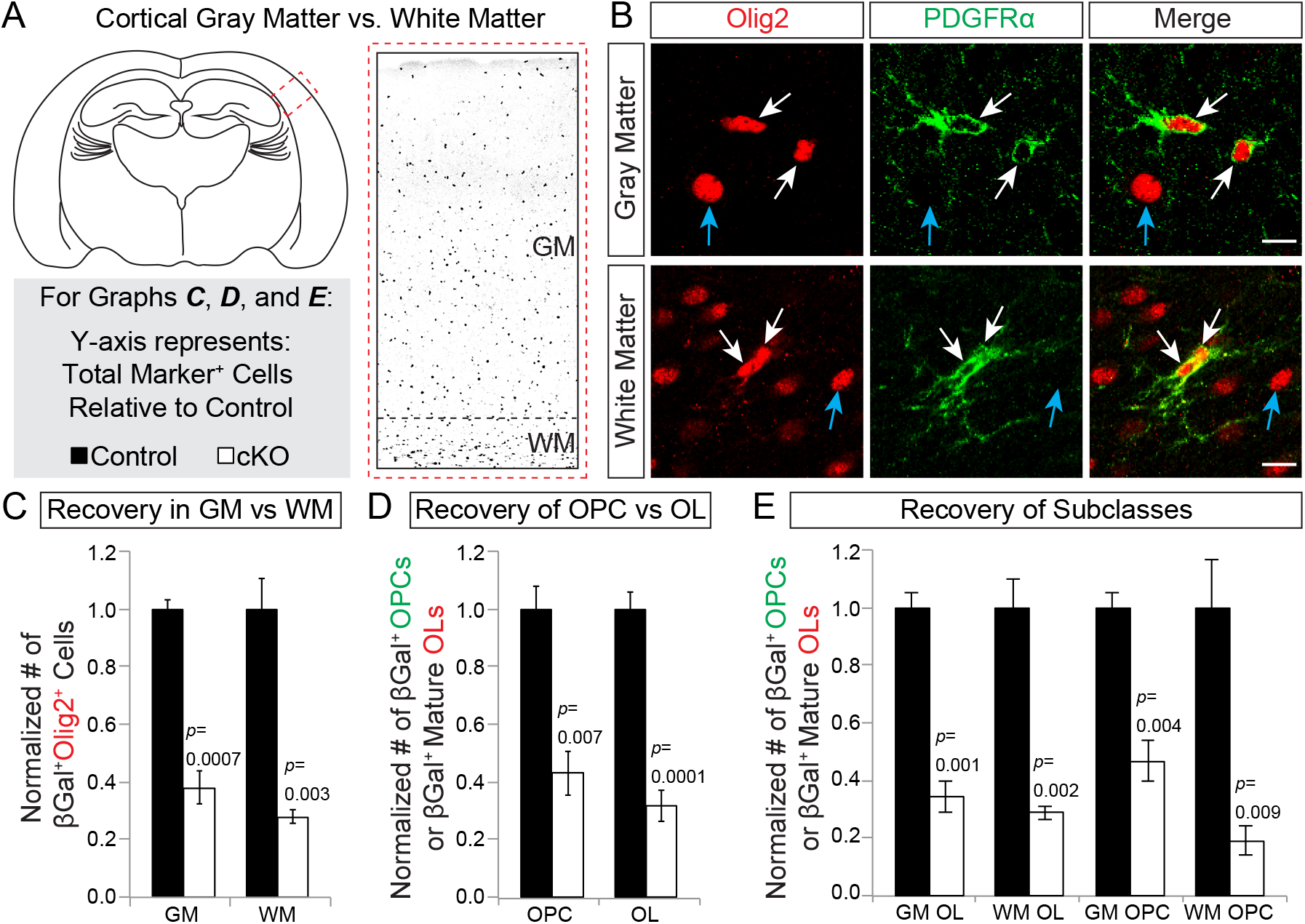
Dorsally derived WM OPCs are the slowest oligodendrocyte lineage population to recover in *Emx1;Smo* mutants at P30. ***A*,** Schematic illustrating the location of the gray matter (GM) and white matter (WM) of the adult somatosensory cortex at P30. Legend for graphs in (C-E). ***B*,** Sections were stained with Olig2 (red) as a marker for the entire oligodendrocyte lineage and PDGFRα (green) as a marker for OPCs. Thus Olig2^+^PDGFRα^+^ cells are OPCs and Olig2^+^PDGFRα^−^ cells are mature oligodendrocytes (OLs). Sections were also stained with βGal to identify dorsally-derived cells (not shown). Representative images of OPCs (white arrows) and OLs (blue arrows) in the GM (top row) and WM (bottom row) of the somatosensory cortex at P30. ***C-E*,** Graphs of the total number of marker^+^ cells in *Emx1;Smo* mutants relative to controls (±SEM among biological replicates). Several distinct populations of dorsally-derived oligodendrocyte lineage cells (βGal^+^Olig2^+^) were analyzed, including GM vs. WM in (C), OPCs (β Gal^+^Olig2^+^PDGFRα^+^) vs. OLs (βGal^+^Olig2^+^PDGFRα^−^) in (D), and a composite analysis of GM OPCs, GM OLs, WM OPCs, and WM OLs in (E). Scale bar, 20 μm.

However, we wondered whether this lack of WM OPCs was an incomplete snapshot of a longer recovery process. During postnatal neocortical development, many OPCs normally differentiate into mature oligodendrocytes. This maturation is especially robust in the white matter, which has been shown to promote oligodendrocyte differentiation (Viganò et al., 2013). Therefore, we reasoned that at P30 we might be analyzing a dynamic WM OPC population that was favoring differentiation over proliferation to help replenish the depleted OL population. To test this idea, we analyzed *Smo* control and cKO brains at two later time points, P90 and P180. Dorsally-derived WM OPCs were indeed slow to recover, reaching only ∼19% of control numbers by P30 and ∼33% that of controls by P90 (Figure 6A). However, by P180 the dorsal WM OPC population had completely recovered (Figure 6A). This recovery was reflected in the total WM OPC population (dorsal plus ventral) as well, which had made a complete recovery by P180 (Fig. 6B). In contrast, dorsally-derived WM OLs only reached a ∼56% recovery compared to controls by P90 and stabilized at that level through P180 (Figure 6C). Interestingly, the total WM OL population had already recovered by P90 (Fig. 6D), suggesting that ventrally-derived cells had compensated for the relative decrease in dorsally-derived WM OLs by this time point. Together these data show that WM OPCs and WM OLs both eventually made a full recovery in the *Smo* cKO neocortex, albeit through different means; WM OPCs reached normal levels by a complete recovery of the dorsally-derived *Smo* mutant population, whereas WM OLs recovered by a combination of dorsal mutants and ventrally-derived wild-type cells.

**Figure 6.**
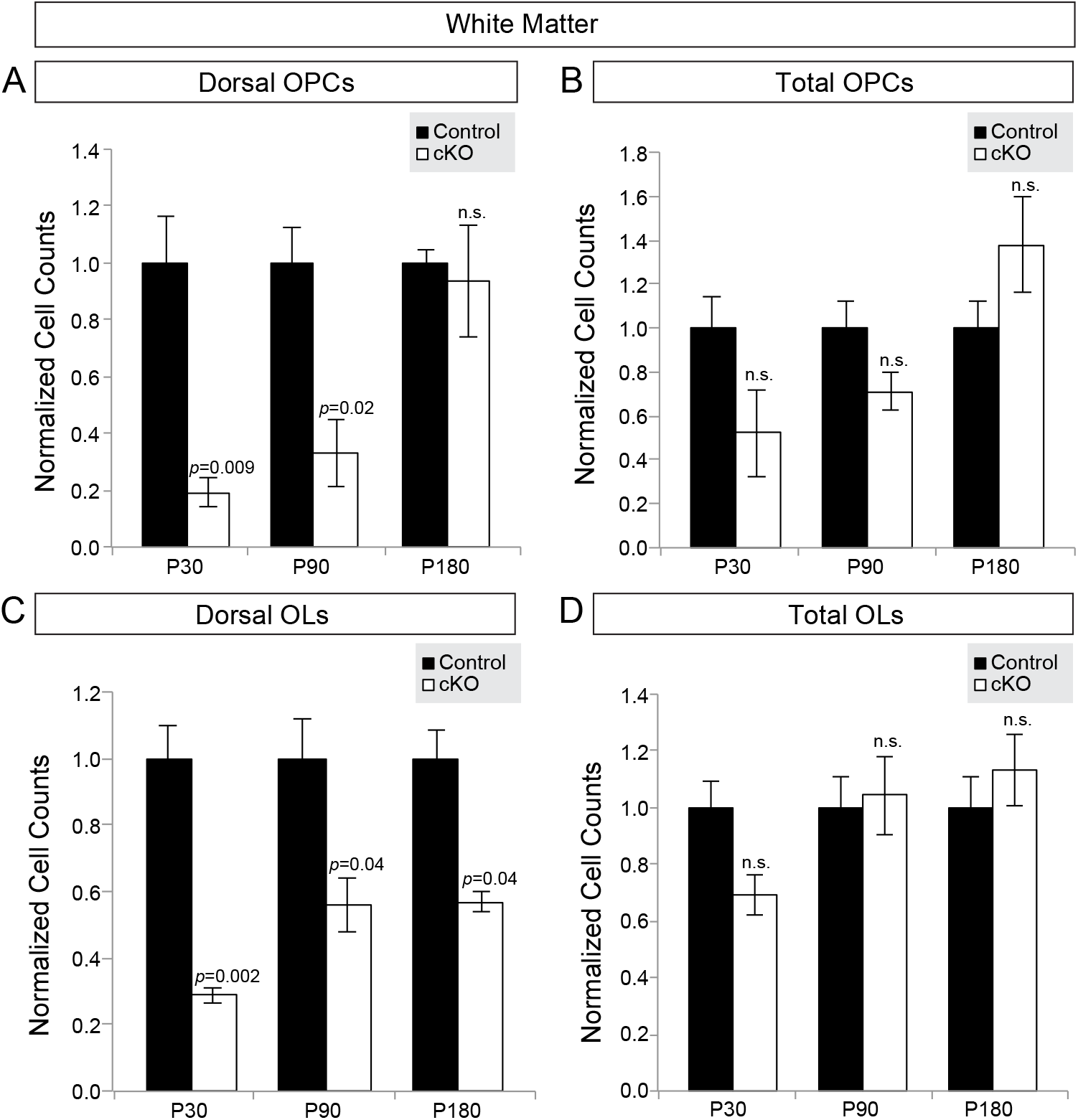
Recovery strategies of OPCs and OLs in the White Matter. For all graphs ***A-D*,** total number of marker^+^ cells in *Emx1;Smo* mutants is relative to age-matched controls (±SEM among biological replicates). ***A, C*,** Dorsally-derived (βGal^+^) OPCs (A) or mature OLs (C) were quantified in the white matter (GM). ***B, D*,** Total (βGal^+^ plus βGal^−^) OPCs (B) or mature OLs (D) were quantified in the white matter (GM). ***A*,** Dorsally-derived WM OPCs (βGal^+^Olig2^+^PDGFRα^+^) fully recover in *Emx1;Smo* mutants following an expansion between P30 and P180. ***B*,** Total WM OPCs (βGal^+^ plus βGal^−^) also completely recover to control levels. ***C*,** Dorsally-derived WM OLs (βGal^+^Olig2^+^PDGFRα^−^) never fully recover by P180 in *Emx1;Smo* neocortex. ***D*,** Total WM OLs (βGal^+^ plus βGal^−^) do make a complete recovery to control levels. Quantifications for P30 re-graphed from Figure 5.

We next analyzed OPCs and OLs in the gray matter of *Smo* control and cKO brains. Just like WM OLs, the dorsally-derived GM OLs never fully recovered to control levels (Fig. 7A), but the total (dorsal plus ventral) GM OL population made a complete recovery between P30 and P90 (Fig. 7B). These data again suggest compensation by ventrally-derived GM OLs. GM OPCs, on the other hand, exhibited a different pattern of recovery. GM OPCs recovered to ∼62% control numbers by P90 and remained stable at this level through P180 (Fig. 7C), despite the total GM OPC population (dorsal plus ventral) never fully recovering to control numbers (Fig. 7D). Therefore, GM OPCs were the only population to not make a complete recovery in the *Smo* cKO neocortex. Taken together, these data indicate several different outcomes as a result of losing Shh signaling in the dorsally-derived oligodendrocyte lineage: 1) WM OPCs made a complete recovery, entirely through expansion of the dorsally-derived *Smo* mutant cells; 2) WM and GM OLs also fully recovered, but did so through dorsal mutant expansion and ventral wild-type compensation; 3) GM OPCs never fully recovered, suggesting impaired expansion of dorsal mutant cells and incomplete compensation by ventral controls cells.

**Figure 7.**
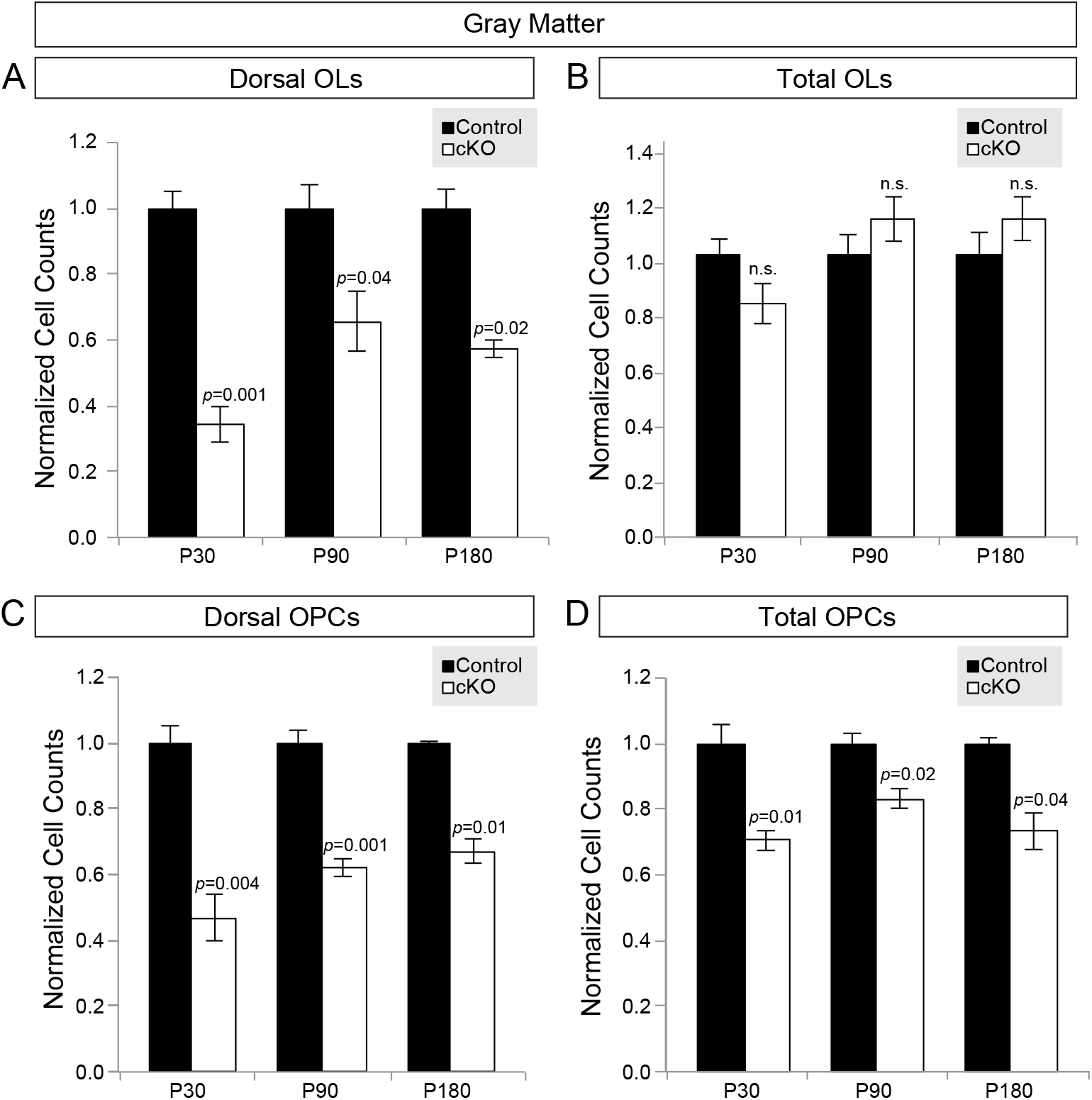
Recovery strategies of OPCs and OLs in the Gray Matter. For all graphs ***A-D*,** total number of marker^+^ cells in *Emx1;Smo* mutants is relative to age-matched controls (±SEM among biological replicates). ***A, C*,** Dorsally-derived (βGal^+^) OPCs (A) or mature OLs (C) were quantified in the gray matter (GM). ***B, D*,** Total (βGal^+^ plus βGal^−^) OPCs (B) or mature OLs (D) were quantified in the gray matter (GM). ***A*,** Dorsally-derived GM OLs (βGal^+^Olig2^+^PDGFRα^−^) do not fully recover by P180 in *Emx1;Smo* mutants. ***B*,** Total GM OLs (βGal^+^ plus βGal^−^) do make a complete recovery to control levels. ***C*,** Dorsally-derived GM OPCs (β Gal^+^Olig2^+^PDGFRα^−^) never fully recover by P180 in *Emx1;Smo* neocortex. ***D*,** Total GM OPCs (βGal^+^ plus β Gal^−^) also never make a complete recovery to control levels. Quantifications for P30 re-graphed from Figure 5.

Finally, we wanted to begin to understand the mechanism underlying the failure of the dorsally-derived GM OPCs to fully recover in the *Emx1;Smo* mutants. One possibility is that GM OPCs are less proliferative in the absence of Shh signaling. To test this hypothesis, we examined the proliferative capacity of dorsally-derived OPCs in *Smo* control and cKO brains at P4, by staining sections for Olig2, βGal and the cell proliferation marker, Ki67 (Fig. 8A). We then quantified the proportion of dorsally-derived Olig2^+^ cells that were Ki67^+^ in *Smo* cKO mutants and littermate controls (Fig. 8A-B). We found that overall the dorsally-derived Olig2^+^ cells in the *Smo* cKO mutants were less proliferative than in the *Smo* wild-type controls (Fig. 8B). Interestingly, this proliferation defect was entirely attributed to GM cells. In the neocortical GM of control animals ∼19% of dorsally-derived Olig2^+^ cells were proliferating, compared to only 12% in *Smo* cKO mutants (Fig. 8C). In contrast, *Smo* cKO mutants and controls had similar proportions of Ki67^+^ WM cells (16% and 15%, respectively). Therefore, in the absence of Shh signaling, dorsally-derived Olig2^+^ cells are less proliferative specifically in the gray matter, but not white matter. This defect may account, at least in part, for the inability of GM OPCs to fully recover in *Smo* cKO mutant brains.

**Figure 8.**
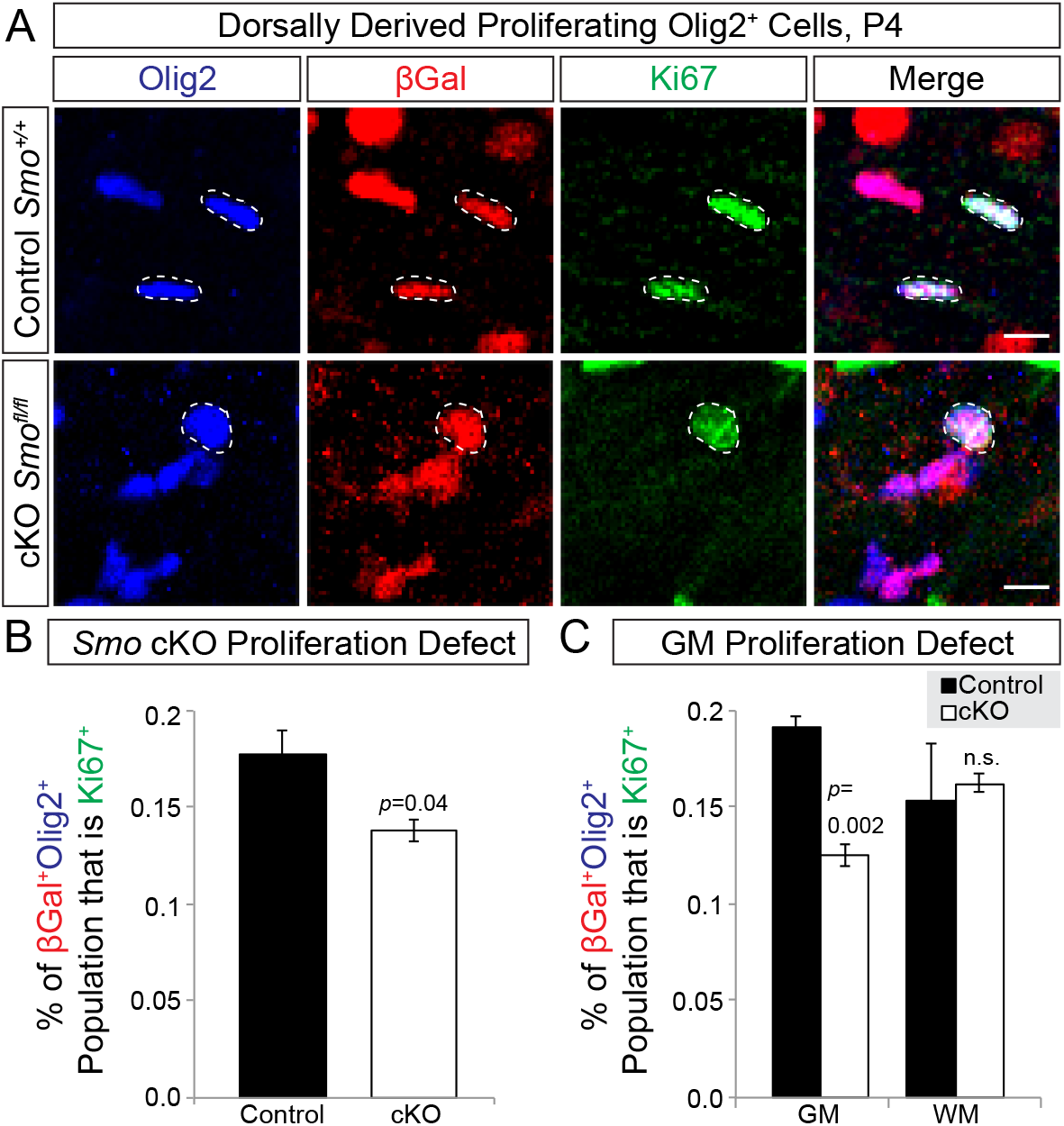
Dorsally derived oligodendrocyte-lineage cells are less proliferative in *Emx1;Smo* mutants compared to controls at P4. ***A*,** Sections were stained with Olig2 (blue) to identify the oligodendrocyte lineage, βGal (red) to identify dorsally-derived cells, and Ki67 (green) to label proliferating cells in *Emx1;S-mo* controls (top row) and mutants (bottom row). Thus, Olig2^+^βGal^+^Ki67^+^ cells represent proliferating cells of the dorsally-derived oligodendrocyte lineage. ***B*,** *Emx1;Smo* mutants exhibited a marked decrease in the percentage of Olig2^+^βGal^+^ cells that were Ki67^+^. ***C*,** The proliferative defect of Olig2^+^βGal^+^ cells was specific to the gray matter (GM) and was not observed in the white matter (WM). Graphs are average percent of total Olig2^+^βGal^+^ cells (±SEM among biological replicates). Scale bars, 10 μm.

## DISCUSSION

We recently showed that Shh signaling to dorsal forebrain progenitors is necessary for normal oligodendrocyte production in the embryonic neocortex (Winkler et al., 2018). Loss of the Shh effector, *Smo*, led to a dramatically reduced oligodendrocyte population at E17.5. In the current study, we employed the same genetic model and followed the subsequent recovery of the remaining oligodendrocyte lineage during postnatal neocortical development. Here, we demonstrate that (1) despite being severely diminished at E17.5, the oligodendrocyte lineage is able to recover robustly in adult *Emx1;Smo* mutants; (2) this recovery is accomplished by a cooperation between the remaining dorsally-derived *Smo* mutant cells and ventrally-derived *Smo* wild-type cells; (3) distinct subpopulations of the oligodendrocyte lineage respond differently during recovery; and (4) dorsally-derived GM OPCs are the subclass most severely affected by loss of Shh signaling, at least partially due to diminished proliferative capacity.

### Contributions of dorsal and ventral sources to oligodendrocyte recovery in the *Smo* mutant neocortex

In the developing mouse forebrain, OPCs first appear in the ventral ventricular zones of the medial ganglionic and lateral ganglionic eminences in two temporal waves, and some of these cells migrate into the dorsal forebrain starting around E15.5 (Kessaris et al., 2005). A third wave of OPC production subsequently starts locally in the dorsal ventricular zone of the neocortex (Kessaris et al., 2005), and as early as E17.5 this dorsal population already makes up the vast majority of oligodendrocyte lineage cells in the neocortex (Winkler et al., 2018). Dorsally-derived cells continue to dominate in the mature neocortex, contributing 80-90% of all oligodendrocytes and OPCs (Tripathi et al., 2011; Winkler et al., 2018). Thus, the neocortical oligodendrocyte lineage comprises a specific ratio of cells that originate from distinct ventral and dorsal forebrain regions. Despite their different origins, however, previous studies have suggested that these different sources of oligodendrocytes are functionally similar and can compensate for each other when one source is lost. For example, genetic ablation of the ventral source did not cause any deficits in oligodendrocyte number or amount of myelin in the corpus callosum, likely due to compensation by dorsally-derived cells (Kessaris et al., 2005). A similar compensation was reported after ablation of the dorsal source, presumably by the unaffected ventrally-derived cells (Kessaris et al., 2005). These studies raised several important questions about oligodendrocyte development in the neocortex. If the different sources of oligodendrocytes are equivalent, what are the mechanisms by which the dorsal population comes to dominate in the neocortex? Are there intrinsic differences that allow the dorsal population to outcompete their ventral counterparts? Or does production of the dorsal lineage simply outpace that of the ventral lineage over time?

Our experimental paradigm in the current study allowed us to address some of these questions. By knocking out the *Smo* gene in Emx1^+^ progenitors, we established a scenario in which the normal ratio of dorsal to ventral OPCs was disrupted. We found that the overall numbers of OPCs and oligodendrocytes eventually recovered in the mature neocortex. This recovery was accomplished in part by a significant expansion of the ventrally-derived oligodendrocyte lineage, thus confirming that during normal development the ventral source is somehow kept in check by the presence of large numbers of dorsally-derived OPCs (Richardson et al., 2006). Interestingly, however, we found that the diminished dorsal population also underwent a substantial expansion postnatally. In fact, the dorsal recovery appeared to outperform the ventral compensation, since the ratio of dorsal to ventral Olig2^+^ cells increased from 1:2 at E17.5 to closer to 1:1 by P30. These results suggest that the competition between dorsal and ventral sources is not simply an issue of starting numbers, but that the dorsally-derived cells hold some sort of advantage over the ventrally-derived cells in the developing neocortex. The cellular and molecular mechanisms that underlie this advantage remain to be elucidated. One idea that has been raised is that the different populations compete for limited resources, such as local factors that promote proliferation or survival (Richardson et al., 2006). Another possibility is that the dorsally-derived oligodendrocyte lineage cells are simply developmentally younger than their earlier-born ventral counterparts, and therefore more likely to proliferate rather than differentiate. Alternatively, dorsally-derived cells may be inherently better at expanding or surviving in the dorsal niche due to intrinsic differences. It will be interesting in the future to test these different possibilities, for example by transplanting equal numbers of developmental age-matched ventral and dorsal OPCs into the neocortex and determining if one source still outperforms the other.

Another interesting feature of the recovery phenotype in *Smo* cKO mutants is that once the total numbers of oligodendrocyte-lineage cells recovered to near wild-type levels, the ratio of dorsal to ventral cells remained fairly steady thereafter. For example, there was substantial overall recovery in the mutants already at P4, and between P4 and P30 (and even at P180, data not shown) the dorsal cells always constituted between ∼45-55% of the total Olig2^+^ population. These data suggest that the dorsally-derived cells only recover as much as is necessary to achieve normal numbers of oligodendrocytes and OPCs when combined with the compensation from the ventral source. Thus, dorsally-derived and ventrally-derived cells in *Emx1;Smo* mutants may strike a functional balance that remains steady once recovery of the total population is sufficient. This could be due to the tight homeostatic control that OPCs exhibit, through a balance of active growth and self-repulsion (Hughes et al., 2013). Future studies should shed more light on this issue, perhaps by testing whether the dorsally-derived oligodendrocyte lineage can recover to a greater extent in the absence of any ventrally-derived cells.

### Oligodendrocyte-lineages cells are heterogeneous in their response to loss of Shh signaling

Another key finding in our study is that distinct subpopulations of the oligodendrocyte lineage responded differently during recovery in *Smo* cKO mutants. We specifically focused on differences between regions (WM vs. GM) and differentiation states (OPC vs. OL). Among the dorsally-derived *Smo* mutant population, WM OPCs performed the best over time and eventually recovered to wild-type numbers. In contrast, recovery was incomplete in the other three populations: GM OPCs, WM OLs and GM OLs. For the WM and GM OL populations, even though the dorsally-derived cells did not reach control numbers, the overall (dorsal plus ventral) populations did fully recovery. In these cases, we cannot determine whether the dorsal mutant OLs stopped expanding because of intrinsic deficiencies, or because they reached an equilibrium with the ventral wild-type cells once the total population fully recovered. For GM OPCs, on the other hand, the overall (dorsal plus ventral) population never reached control numbers, yet the dorsally-derived mutant cells only recovered to 67% of control levels. Therefore, the *Smo* mutant OPCs in the GM appear to have reached some cell-autonomous upper limit on their recovery. One possible explanation for these observations is that the GM OPCs depend more on Shh signaling for expansion, consistent with the known mitogenic effects of Shh on many different cell types.

WM OPCs, on the other hand, may represent a population of the oligodendrocyte lineage that is Shh-independent. Several studies have provided evidence for a Shh-independent pathway for oligodendrogenesis in the dorsal spinal cord, possibly via an FGF-dependent signaling mechanism (Cai et al., 2005; Chandran et al., 2003; Fogarty et al., 2005; Vallstedt et al., 2005). Thus, a similar Shh-independent population of oligodendrocyte lineage cells could exist in the dorsal forebrain, which may explain the ability of *Smo* mutant WM OPCs to fully recover. Interestingly, a recent study identified a population of oligodendrocyte-lineage cells at the subcallosal ventricular-subventricular zone that is Shh dependent within the first postnatal week in mice (Tong et al., 2015). Genetic ablation of *Smo* in dorsal ventricular-subventricular zone progenitors at P0 resulted in reduced production of oligodendroglial cells in the corpus callosum at P28 (Tong et al., 2015). Consistent with these data, we find that at P30 the *Smo* mutant Olig2^+^ cells as a whole are significantly reduced. The fact that the mutant WM OPCs eventually make a full recovery by P180, but mutant WM OLs do not, might indicate a specific function of Shh in OL differentiation. It would be interesting to determine if the Shh-dependent population identified by Tong et al. primarily comprises mature OLs. Such a result could indicate that the initial generation of OPCs from the postnatal dorsal ventricular-subventricular zone is Shh-independent, whereas their subsequent differentiation into mature OLs requires Shh signaling.

It is also important to consider how the local niches might affect recovery in the *Smo* cKO mutant neocortex. Previous studies have shown that GM OPCs have a longer cell cycle length and lower differentiation properties than WM OPCs (Dimou et al., 2008; Kang et al., 2010; Young et al., 2013). Furthermore, WM OPCs respond to PDGF, but GM OPCs do not (Hill et al., 2013). Additionally, in a series of homotopic and heterotopic transplants in adult mice, OPCs derived from the GM could partially differentiate in the WM, but not in the GM (Viganò et al., 2013). Together, these studies suggest certain differences in specific niche environments, and raise the possibility that one reason WM OPCs are able to recover so well is at least partially due to the fact that they inhabit a more permissive and supportive environment than GM OPCs.

Besides a potential role for Shh in differentially regulating proliferation and/or maturation in distinct Olig2^+^ populations, recent evidence indicates that differing levels of Shh signal are sufficient to drive different cell fates within the oligodendrocyte lineage. In the zebrafish spinal cord, some OPCs rapidly differentiate into OLs while others remain as NG2^+^ OPCs (Ravanelli et al., 2018). These two populations arise sequentially from distinct neural progenitors and, thus, represent distinct lineages (Ravanelli et al., 2018). Importantly, these lineages are specified by small differences in Shh signaling, where higher levels resulted in an OL fate and lower levels of Shh signaling resulted in an NG2^+^ OPC fate (Ravanelli et al., 2018). If a similar mechanism is at work in the developing forebrain, this could account for some of the differential effects we see on the subclasses of neocortical Olig2^+^ cells. Indeed, at P180 we find that OLs are more severely affected by *Smo* knockout than their OPC counterparts within the same region (WM or GM). Interestingly, a number of recent studies have started to reveal diversity within the oligodendrocyte lineage, demonstrating that oligodendrocytes are not a homogeneous population (for recent reviews, see (Akkermann et al., 2017; Dimou and Simons, 2017; Newville et al., 2017)). Different oligodendrocyte subpopulations can be distinguished based on morphology (Chong et al., 2012; Osanai et al., 2017), locations in the white matter or gray matter (Dimou et al., 2008; Viganò et al., 2013; Vue et al., 2014), gene expression profiles (Butt et al., 1998b; Kleopa et al., 2004; Marques et al., 2016; Nakatani et al., 2013), molecular markers (Boda et al., 2011; Chen et al., 2009; Fumagalli et al., 2011; Kukley et al., 2007; Ziskin et al., 2007), proliferation and differentiation potential (Dimou et al., 2008; Hughes et al., 2013; Kang et al., 2010; Young et al., 2013), and their myelination properties (Anderson et al., 1999; Battefeld et al., 2016; Bjartmar et al., 1994; Butt et al., 1998a; Crawford et al., 2016; Tomassy et al., 2014). It remains to be determined which of these properties reflect distinct cell lineages and which are simply different maturational states or functions of external influences from their environments. Future studies using single-cell RNA sequencing approaches and clonal analysis through genetic fate-mapping should shed light on this emerging topic. It will also be important to combine these approaches with loss- and gain-of-function for Shh signaling to determine the precise roles that this pathway plays in driving oligodendrocyte diversity.

## ACKNOWLEDGEMENTS

This work was supported by the Children’s Hospital Colorado Program in Pediatric Stem Cell Biology (S.J.F.), The Boettcher Foundation (S.J.F.), and NIH/NINDS R56NS109239-01 (S.J.F.). The authors declare no competing financial interests.

